# Low-dose fluvoxamine modulates endocytic trafficking of SARS-CoV-2 spike protein: a potential mechanism for anti-COVID-19 protection by antidepressants

**DOI:** 10.1101/2021.06.15.448391

**Authors:** Oleg O Glebov

## Abstract

Commonly prescribed antidepressants may be associated with protection against severe COVID-19, with one drug (fluvoxamine) currently undergoing a Phase 3 clinical trial. The mechanism of their action, however, remains unknown. Here, I investigated the effect of fluvoxamine on membrane trafficking of the SARS-CoV-2 spike protein and its cell host receptor ACE2 in HEK293T cells. A sub-therapeutic concentration (80 nM) of fluvoxamine rapidly upregulated fluid-phase endocytosis, resulting in enhanced accumulation of the spike-ACE2 complex in enlarged early endosomes. Diversion of endosomal trafficking may provide a simple cell biological mechanism consistent with the protective effect of antidepressants against COVID-19, highlighting their therapeutic and prophylactic potential.

## Main text

More than a year of the COVID-19 pandemic has brought about ongoing disruption of daily activities, severe economic downturn and unprecedented healthcare crisis across most of the world’s nations. Although relatively rapid COVID-19 vaccine development has been generally considered a success, vaccine availability is currently limited by the logistical difficulties pertaining to production, storage and distribution^1^; furthermore, periodic emergence of novel strains of the SARS-CoV-2 virus fuels recurring concerns about vaccine efficacy and future-proofing^2^. Therefore, repurposing of cheap, readily available and safe drugs against COVID-19 remains a major healthcare priority.

One promising group of candidate drugs is antidepressants, which have been linked to protection against severe COVID-19 in several post-hoc clinical studies^3^, and may block SARS-CoV-2 infection and replication in cell models^4–7^. Foremost amongst these is a generic selective serotonin reuptake inhibitor (SSRI) fluvoxamine, that has been shown to protect against severe COVID-19 in a Phase 2 trial^8^ as well as a real-world study^9^ and is currently undergoing a Phase 3 study^10^. The suggested protective effect of fluvoxamine has been tentatively linked to modulation of immune response through activation of the ER stress and sigma receptor^11^, or blockade of acid sphingomyelinase^7,12,13^, however the underlying cell biological mechanism remains unknown.

It had been suggested that pharmacological modulation of SARS-CoV-2 membrane trafficking might offer a strategy for both treatment and prophylaxis of COVID-19, and a list of candidate drugs included fluvoxamine^14^. Given the emergence of clinical data suggesting efficacy of fluvoxamine against COVID-19, the effect of fluvoxamine treatment on membrane trafficking was assessed in a cell line. To this end, HEK293T cells were treated with a range of fluvoxamine concentrations. Although standard dosing of fluvoxamine results in considerable brain enrichment of the drug (10 uM)^15^, its concentrations in the blood plasma are much lower^16^, therefore a range of concentrations was used. Treatment for 1 h resulted in a significant increase in endocytosis, as measured by the membrane FM dye uptake (**Figure 1a,b**). Remarkably, this increase was apparent at as little as 80 nM fluvoxamine, equivalent to approximately 1/5th of the blood plasma concentration^16^.

**Figure 1.**
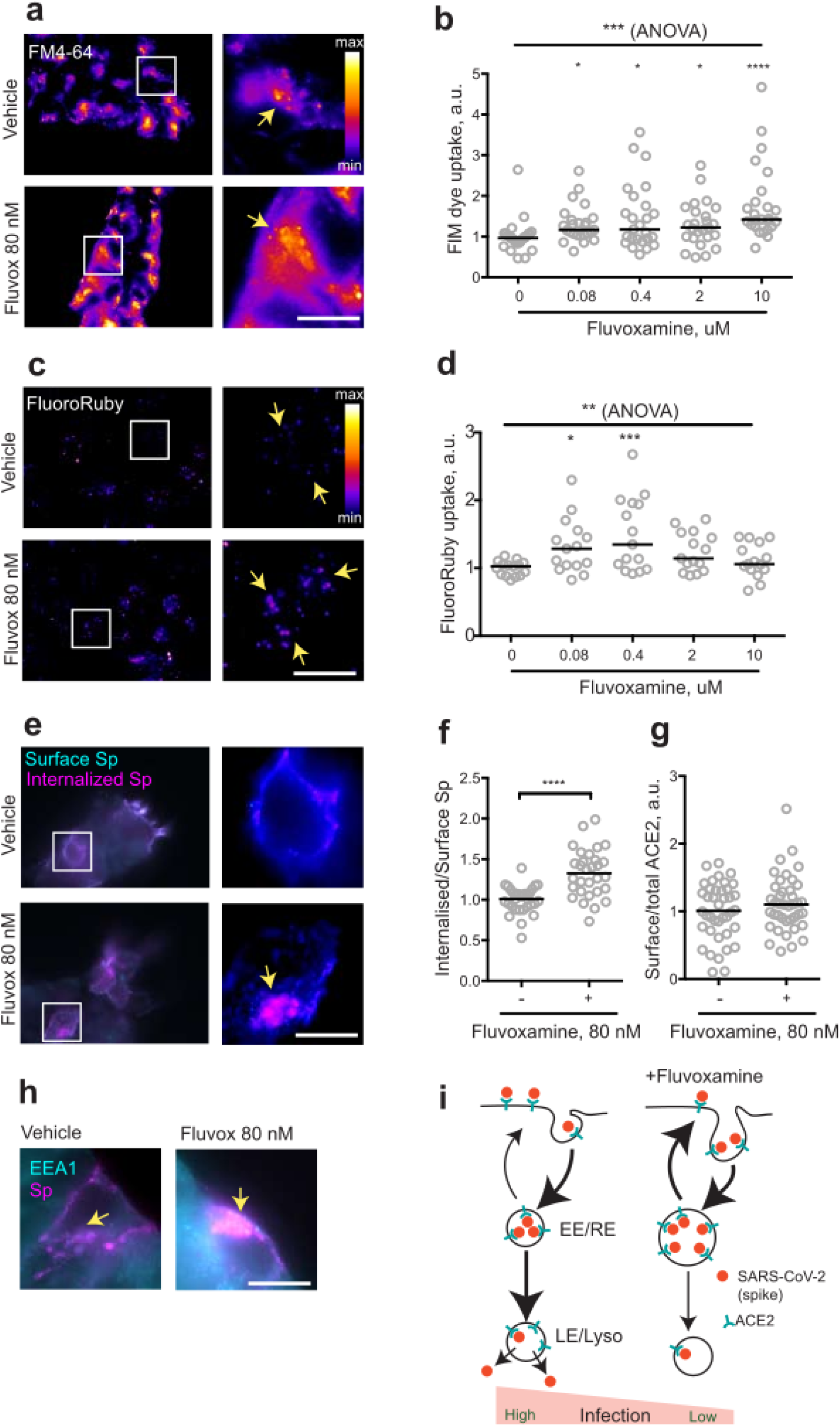
Fluvoxamine modulates endocytosis, including that of SARS-CoV-2 Spike. **a,** HEK293T cells were allowed to internalise FM dye for 1 hour. Representative images of cells incubated with vehicle or 80 nM Fluvoxamine. Arrows denote puncta of internalised label. **b**, Quantification of FM dye uptake experiments for various concentrations of fluvoxamine. ***P=0.0001, Kruskal-Wallis test. ****P<0.0001, *P<0.05, Dunn’s multiple comparisons test. **c**, HEK293T cells were allowed to internalise FluoroRuby for 1 hour. Representative images of cells incubated with vehicle or 80 nM Fluvoxamine. Arrows denote puncta of internalised label. **d**, Quantification of FluoroRuby uptake experiments for various concentrations of fluvoxamine. ***P<0.001, 1-way ANOVA; ***P<0.001, *P<0.05, Holm-Sidak’s multiple comparisons test. **e,** HEK293T cells were transfected with a plasmid expressing human ACE2, incubated with the recombinant SARS-CoV-2 spike protein for 1 hour and sequentially immunostained for surface (cyan) and internalised (magenta) spike protein. Representative images of cells incubated with vehicle or 80 nM Fluvoxamine. Arrows denote apparent puncta of internalised spike protein. **f**, Quantification of the spike protein uptake assay. ****P<0.0001, unpaired two-tailed t test. **g**, Quantification of surface ACE2 (live-labelled by spike protein) lvs total ACE2 labelling following 1h treatment with 80nM fluvoxamine. P>0.05, Mann Whitney test. **h**, Accumulation of Spike in EEA1-positive early endosomes following 1 hour treatment with vehicle or 80 nM fluvoxamine. Arrows denote apparent puncta of internalised spike protein. **i**, Proposed model for modulation of SARS-CoV-2 endocytic trafficking by fluvoxamine. Fluvoxamine promotes trafficking of SARS-CoV-2 through early/recycling endosomes (EE/RE), diverting it away from the infectious step involving late endosomes/lysosomes (LE/Lyso). Thickness of the arrows represents relative intensity of trafficking. Scale bar, 10 um.

To verify upregulation of endocytosis, uptake of the fluid-phase biologically inert endocytic marker fluorescent dextran FluoroRuby was also measured. Treatment with low concentrations of fluvoxamine resulted in a significant increase in FluoroRuby uptake mirroring the effect on the FM dye uptake, confirming the increase in both the membrane area and the volume of endocytosis (**Figure 1c,d**). Higher concentrations of fluvoxamine resulted in cell rounding and detachment (data not shown), indicating a possible cytotoxic effect. Taken together, these observations suggest that sub-therapeutic doses of fluvoxamine may rapidly alter endocytosis in a non-specific manner.

The non-specific nature of the fluvoxamine effect suggested that it could also affect trafficking of SARS-CoV-2 as postulated previously^14^. To investigate this possibility, the key steps of SARS-CoV-2 ingress were recreated by imaging internalisation of recombinant SARS-CoV-2 spike protein^17^ into HEK293T cells transiently transfected with a plasmid expressing the human SARS-CoV-2 receptor ACE2^18^. Consistent with the effect on fluid-phase and membrane endocytosis, 1 h incubation with 80 nM fluvoxamine resulted in a significant increase in spike protein internalization, as measured by the ratio of the internal spike labelling to that at the surface (**Figure 1e,f**). Relative levels of surface-bound spike protein were not changed following 1 h treatment, suggesting that surface binding to the ACE2 and recycling of the internalised ACE2 back to the plasma membrane were unaffected by the drug treatment (**Figure 1g**). From these results it can be concluded that fluvoxamine treatment upregulates endocytosis of spike protein.

Following endocytosis, internalised cargoes proceed to early endosomes, where they undergo sorting to be recycled back to the cell surface or trafficked into late endosomes/lysosomes. Transit through early endosomes has been implicated as a key stage in coronaviral infection^12,19–22^. To investigate the effect of fluvoxamine treatment on early endosomes, cells were immunostained for a canonical early endosome marker EEA1. In line with previously published data^19^, internalised spike protein co-localised with EEA1-positive puncta, suggesting that early/recycling endosomes are likely to be involved in SARS-CoV-2 trafficking (**Figure 1h**). Fluvoxamine treatment significantly enlarged EEA1 positive puncta, consistent with increased endosomal capacity as evidenced by the increase in fluid-phase uptake (**Figure S1a**); in contrast, the morphology of late endosomes as shown by immunostaining for a late endosome marker LAMP1 was not visibly affected. Thus, low doses of fluvoxamine may enhance trafficking through early but not late endosomes.

Two more antidepressants identified as candidates for modulating SARS-CoV-2 membrane trafficking^14^ were a SSRI sertraline and a tricyclic drug imipramine, both of which have recently been shown to block SARS-CoV-2 infection in epithelial cells^7^. Their effect on endocytosis was tested as described above. Treatment with low therapeutic concentrations of sertraline (80 nM) and imipramine (200 nM) resulted in an increase in endocytosis as evidenced by FM dye uptake (**Figure S1b,c**). Higher concentrations of sertraline did not produce an increase in FM uptake, consistent with either cytotoxicity or previously reported inhibition of dynamin function and endocytosis blockade^23^. Global upregulation of endocytosis may therefore represent a generalized feature of low-dose antidepressant treatment.

In summary, it was found that a short-duration treatment with a sub-therapeutic concentration of fluvoxamine results in a significant rearrangement of membrane trafficking in a human cell line, enhancing endocytic uptake of SARS-CoV-2 spike proteins through early endosomes. This finding is consistent with a mechanism involving re-routing of the internalised ACE2/Spike complex into the early/recycling endosome away from the late endosome/lysosome (**Figure 1i**). Given the emerging role of the late endosome/lysosome in both SARS-CoV-2 infection and release^12,20,24–28^, preferential retention in the early/recycling endosome may provide a cell biological framework underlying the putative clinical benefits of fluvoxamine. The molecular mechanisms behind this cell biological process, however, remain to be determined. The sigma receptor-independent mode of action is supported by the similarity in the effects of low concentration fluvoxamine (sigma receptor agonist) and sertraline (sigma receptor antagonist). At micromolar concentrations, fluvoxamine, sertraline, imipramine and other antidepressants and antipsychotics may act as functional inhibitors of acid sphingomyelinase (FIASMA)^29^; another noteworthy possibility is direct regulation of the plasma membrane properties through lipophilic association with the lipid bilayer^30^.

Only the drugs from the previously published list of candidates^14^ were included in this work, with most of the focus being on fluvoxamine due to its clinical potential. However, emerging laboratory and clinical evidence hints at the possible therapeutic potential of other antidepressants^3–5,7,12^. Diversion of SARS-CoV-2 away from its productive infectious route could imply that antidepressants may not only protect against severe COVID-19, but prevent SARS-CoV-2 infection in the first place. Given the wide prevalence of SSRI prescriptions in outpatient treatment of moderate psychiatric disorders, the prophylactic potential for SSRIs may be assessed by performing post-hoc studies of COVID-19 incidence in the general population.

It must be emphasised that, given the ongoing proliferation of non-peer-reviewed COVID-19 publications^31^, the findings reported here should be interpreted with caution and chiefly considered as a basis for further investigation. Importantly, antidepressant-dependent SARS-CoV-2 membrane trafficking needs to be tracked in more detail and verified in physiologically relevant systems featuring infection with live SARS-CoV-2 particles^32^. Finally, the reported impact on membrane trafficking mechanisms may be of relevance in the context of multiple side-effects associated with antidepressants, suggesting that cell biology of psychiatric drugs outside the central nervous system^33^ merits further investigation.

## Materials and Methods

### Materials

Cell culture materials were from Gibco. Drugs were from Sigma-Aldrich. FM Dyes and FluoroRuby were from Thermo Fisher. Recombinant SARS-CoV-2 spike protein and the anti-spike mouse polyclonal antibody were kindly provided by Dr. Paul F. McKay (Department of Infectious Disease, Faculty of Medicine, Imperial College London) through the COVID-19 Crowdfight initiative. The following rabbit antibodies were used: anti-EEA1, anti-ACE2 (GeneTex), anti-LAMP1 FITC conjugated (Sino Biological), anti-6xHis HRP conjugated (Abcam). Secondary antibodies were from Jackson Immunoresearch.

### Cell culture and uptake assays

HEK293T cells were cultured in DMEM supplemented with 10% bovine serum. For experiments, cells were grown on 13mm round thickness 1.5 coverslips coated with poly-L-Lysine. Cells were transfected using Lipofectamine 2000 and used 24-48 hours afterwards. All uptake experiments were performed at 37°C. For uptake experiments, probes were diluted in culture medium and added to the coverslips in presence of indicated concentrations of drugs or vehicle (DMSO) for indicated periods of time. FM dyes (FM4-64fx and FM1-43fx) were used at the final concentration of 20 ug/ml, FluoroRuby was used at the final concentration of 0.1% w/v, Spike protein was used at the final concentration of 0.14 ug/ml. For FM dye and FluoroRuby uptake measurements, cells were fixed, mounted and imaged without further processing.

### Immunocytochemistry and microscopy

All steps were performed at room temperature. Cells were fixed for 10 min in 4% w/v paraformaldehyde in PBS, permeabilized in the ICC buffer (0.1% w/v Triton X-100, 5% w/v horse serum in PBS), stained with the primary antibodies for 1 hour. The following antibody dilutions (v/v) were used: 1/1000 (anti-spike), 1/1000 (anti-6xHis), 1/500 (anti-ACE2), 1/400 (anti-EEA1), 1/400 (anti-LAMP1). For spike protein uptake measurements, cells were fixed, immunostained with the anti-6xHis antibody for 30 min to label the surface pool, permeabilized, immunostained with an anti-spike antibody for 30 min to label the total spike pool. For surface and total ACE2 measurements, coverslips were live-labelled with spike protein for 30 min before fixation to label the surface pool of ACE2. Following fixation and permeabilization, coverslips were labelled with the anti-ACE2 antibody to visualise the total pool of ACE2. Finally, coverslips were incubated with the anti-rabbit-Alexa Fluor 488 and anti-mouse Alexa Fluor 594 secondary antibodies for 30 min. Coverslips were mounted in Fluoromount-G medium (Southern) and imaged in the EVOS M7000 widefield imaging system (Thermo Fisher) using the 100x objective. Quantification of the images was carried out in ImageJ. For measurement of Spike internalisation, intensity of total Spike labelling was divided by intensity of the surface Spike labelling. All data was normalised to the median values in the vehicle-treated sample.

### Statistical analysis

Datasets represent results of at least 3 independent experiments. Statistical analysis was carried out using the Prism 6.0c software package (GraphPad Software). Data distributions were assessed for normality. For normally distributed datasets, Student’s t-test, 1-way ANOVA and Holm-Šidák’s post-test were used; otherwise, Mann-Whitney rank test, Kruskal-Wallis test and Dunn’s post-test were used. Datasets were presented as scatter dot plots with line at median or as cumulative probability plots.

## Acknowledgements

The author is grateful to Dr Paul F. McKay and Dr Georgina H. Cornish for sharing materials, and to the COVID-19 Crowdfight initiative (https://crowdfight.org) for assistance. EVOS M7000 imaging system was kindly provided by Dr Richard Killick. O.O.G. is supported by The Lewy Body Society (OOG2019/2020) and National Natural Science Foundation of China (32070772).

---

**Figure S1.**
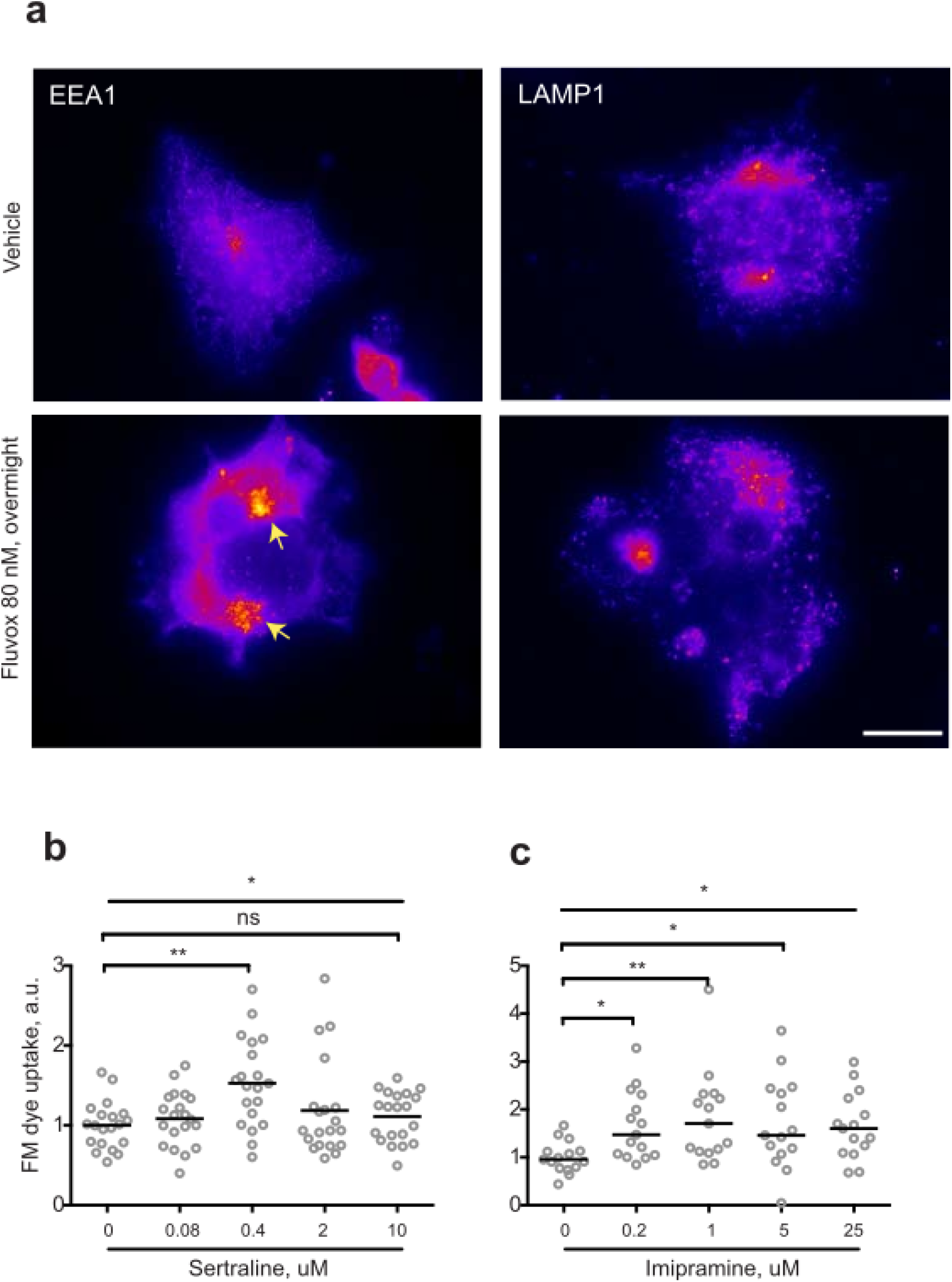
Supplementary data. **a**, HEK293T cells were treated with vehicle or 80 nM fluvoxamine overnight and immunostained for EEA1 or LAMP1. Arrows denote enlarged early endosomes following fluvoxamine treatment. **b**, Quantification of FM dye uptake experiments for various concentrations of sertraline. *P<0.05, Kruskal-Wallis test. **P<0.01, Dunn’s multiple comparisons test. **c**, Quantification of FM dye uptake experiments for various concentrations of imipramine. *P<0.05, Kruskal-Wallis test. *P<0.05, **P<0.01, Dunn’s multiple comparisons test. Scale bar, 20 um.

